# SynBioCrow: provenance-preserving integration of complementary retrosynthesis engines for biosynthetic pathway design

**DOI:** 10.64898/2026.10.06.756997

**Authors:** Thomas J. Heiman

## Abstract

Motivation: Retrobiosynthesis tools use different reaction rules, search strategies and biochemical priors, so each can recover transformations missed by others. Integrating their outputs could broaden pathway coverage, but ensembling can also worsen ranking. Results: SynBioCrow runs independent retrosynthesis engines, normalizes predictions into a provenance-preserving reaction graph, and separates route generation from biochemical evidence, sequence design and bounded learning. On the frozen 65-path Galaxy-SynBioCAD literature benchmark used for SynBioCrow 2.3, connectivity-level partial recovery was 19/65 for the ensemble, 13/65 for RetroBioCat2, 9/65 for RetroPath and 0/65 for DORAnet; the ensemble preserved two exact RetroPath routes but worsened their ranks. SynBioCrow 2.4 subsequently repaired structured-reaction parsing and target-connected Candidate generation and prospectively locked two new external targets, 1,3-propanediol and 1,4-butanediol, before pathway truth was examined. RetroBioCat2 produced 3 and 2 backend-internal SOLVED search outcomes, respectively, whereas DORAnet produced none; however, neither backend recovered any predeclared explicit literature intermediate by exact structure. The negative prospective result was frozen without post-reveal tuning. Production ranking in 2.4 is therefore NONE and the release remains Candidate-stage; validated complete-pathway recovery is not claimed. Availability: SynBioCrow 2.4.0 is archived at DOI 10.5281/zenodo.23173173 with source code and frozen supporting evidence.

## 1 Introduction

Retrobiosynthesis searches backward from a target molecule toward compounds available from a host or defined precursor set. Generalized biochemical transformations can expose non-native production routes, but iterative rule application rapidly expands the search space. Consequently, reaction-rule scope, compound normalization, pruning, search depth and ranking strongly influence which pathways are recovered.

Existing systems expose different parts of this design space. RetroPath2.0 performs rule-based metabolic retrosynthesis [1], with RetroRules providing stereochemistry-aware generalized biochemical reaction rules used by retrosynthesis workflows [10]; RetroBioCat focuses on biocatalytic reaction knowledge and enzyme precedent [2]; DORAnet supports configurable reaction-network expansion [11]; and BioPKS Pipeline uses RetroTide for chimeric type I polyketide-synthase design together with DORAnet for post-PKS enzymatic transformations [12]. Galaxy-SynBioCAD demonstrates how retrosynthesis, pathway scoring and genetic design can be connected in an automated synthetic-biology workflow [3]. These systems are complementary rather than interchangeable.

An earlier motivation for this architecture was the accessibility goal exemplified by Cameo, a high-level Python framework designed to make in silico cell-factory design available through a unified interface to both experienced modelers and users new to metabolic modeling [13]. Cameo demonstrated the value of hiding heterogeneous computational methods behind a simpler design workflow, including enumeration of native and heterologous production pathways. SynBioCrow extends this usability and integration motivation to a contemporary retrosynthesis ecosystem in which useful capabilities are distributed across independently developed engines with different reaction representations, dependencies, interfaces and biochemical scope. Its modular backend boundary is intended to reduce dependence on any single software stack: generators can be updated, replaced or allowed to fail independently while normalized routes and provenance remain available to the downstream workflow.

ChemCrow, an LLM-powered chemistry agent that coordinates specialized computational tools for chemical problem solving, provided a further conceptual inspiration for SynBioCrow and directly influenced the project name [14]. SynBioCrow adapts the broader tool-orchestration idea to computational biosynthetic pathway design, while its evaluated architecture and claims are specific to the provenance-preserving multi-engine workflow described here.

SynBioCrow tests whether independent engines can be treated as complementary proposal sources rather than forcing a single generator to define the reachable reaction space. Predictions are normalized into a shared reaction graph while retaining edge-level provenance. This gives a precise ensemble hypothesis: graph union should preserve constituent routes and may broaden reaction recovery, but improved exact-route retrieval is not assumed and must be demonstrated empirically.

A second design principle is separation of proposal from evidence. Candidate reactions and routes are not promoted simply because a generator produced them. Evidence, enzyme precedent, thermodynamic information, sequence provenance and construct quality are evaluated downstream, with unresolved information represented as abstention rather than positive evidence.

Here we evaluate the frozen SynBioCrow 2.3 baseline using a breadth panel, a route-preservation audit, a frozen held-out literature benchmark, a 2D-versus-3D similarity ablation and computational Design-Build-Test-Learn (DBTL) demonstrations, and then report a prospectively locked external evaluation of the repaired SynBioCrow 2.4 Candidate-generation architecture. The 2.3 benchmark is retained as historical evidence and was not reused to tune 2.4. The central question is not whether an ensemble produces more routes, but whether integration preserves validated constituent chemistry, broadens literature-reaction recovery and supports prospective recovery on new targets without post-truth tuning. The principal contributions are: (i) a modular multi-engine execution layer spanning complementary rule-based, biocatalytic and PKS-oriented generators; (ii) provenance-preserving normalization and reaction-graph union that retain constituent solutions; (iii) fail-closed evidence and computational DBTL handling; (iv) a sealed, frozen 2.3 held-out evaluation that separates reaction coverage from exact-route ranking; and (v) a prospectively locked 2.4 external evaluation whose negative result is preserved as part of the scientific record.

## 2 Methods

### 2.1 System architecture

The frozen SynBioCrow 2.3 baseline integrates DORAnet [11], RetroBioCat2 (the current RetroBioCat implementation lineage) [2], a KNIME-free RetroPath-compatible runtime using RetroPath/RetroRules concepts [1,10], and a BioPKS/RetroTide bridge [12]. Each backend executes independently and emits Candidate records with backend provenance. Molecular identities and reactions are normalized before graph union; duplicate or equivalent edges can retain multiple backend sources. Route enumeration operates on the union graph, permitting a route to contain edges proposed by different engines. SynBioCrow 2.4 retains this proposal/evidence separation while repairing structured-reaction parsing and target-connected Candidate-chain reconstruction for prospective evaluation.

**Figure 1.**
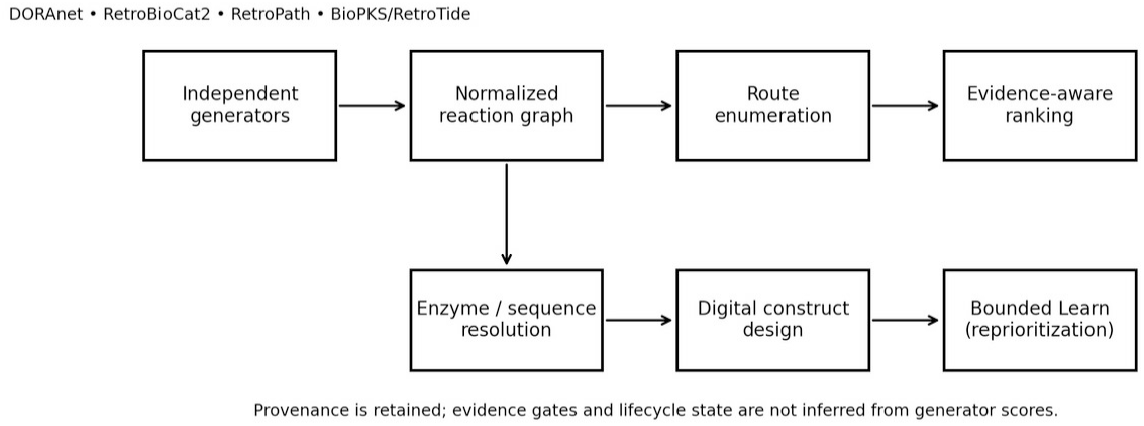
**SynBioCrow computational workflow. Independent generators feed a provenance-preserving normalized reaction graph. Route generation is separated from evidence evaluation, enzyme/sequence resolution, digital construct design and bounded reprioritization.** *Alt text: Workflow diagram showing independent retrosynthesis generators feeding a normalized, provenance-preserving reaction graph, followed by route enumeration, evidence evaluation, sequence-backed construct design and bounded learning*.

### 2.2 Route preservation and breadth benchmark

A 24-target breadth panel characterized candidate coverage, complete-route coverage, route diversity and backend failure modes. For every target with a complete constituent-backend route, persisted candidate checkpoints were audited at the normalized reaction-edge level. The audit distinguished loss of an edge after graph union from failure to return an intact route within a bounded enumeration cap. No backend was rerun for this audit.

### 2.3 Literature benchmark and holdout policy

The main biological benchmark used the Galaxy-SynBioCAD literature pathway collection [3]. The benchmark tests recovery of reported literature chemistry rather than experimental superiority of alternative routes. A deterministic 12-pathway development subset was used for similarity-policy selection and ranking diagnostics; the remaining 65 pathways formed the frozen held-out denominator. One record, 3-methylbutanol (literature_10), was mapping-limited because no usable normalized target structure was available; it remained in the denominator, leaving 64 runnable targets. Because constituent backends differ in intended domain, rule sets, and search behavior, comparisons are interpreted as matched performance within the SynBioCrow execution/evaluation contract rather than as universal rankings of the underlying software systems.

Predictions were generated without access to benchmark reaction truth and SHA-256 sealed before scoring. Evaluation used strict stereochemical reaction identity and connectivity-level matching with stereochemistry removed. Metrics included partial pathway recovery, best reaction recall, exact-route rank, Top-K recovery, mean reciprocal rank (MRR), runtime failures and mapping limitations. Held-out truth was not used for learning, post-holdout ranking revision or policy selection. Results are reported primarily as descriptive benchmark counts and ranks; no inferential significance test is claimed for differences between backends because targets are heterogeneous, backend domains differ, and the frozen benchmark was designed as an engineering evaluation rather than a population sample.

### 2.4 Similarity-policy ablation

The development study compared Morgan/Tanimoto 2D-only ranking [8] with USRCAT 3D-only and USRCAT 3D-first with explicit 2D fallback [7]. Candidate routes were held fixed so that only ranking policy changed. Leave-one-pathway-out evidence retrieval prevented a pathway from supplying its own literature reactions during ranking diagnostics.

### 2.5 Evidence-aware DBTL

Design comprises multi-backend pathway generation, graph integration, route enumeration and ranking. Build resolves provenance-backed proteins and coding sequences, including reviewed UniProt records [5], performs translation-preserving codon design and sequence QC, and assembles regulatory cassettes only when verified part sequences are available. Test evaluates route closure, biochemical-database support including Rhea [4], enzyme evidence, thermodynamics, reproducibility and construct QC. Learn can update bounded prioritization weights from structured TestOutcome records, but cannot create evidence, change evidence-gate outcomes, access protected benchmark truth or promote lifecycle state.

### 2.6 Prospective external evaluation of SynBioCrow 2.4

SynBioCrow 2.4 was evaluated only after the 2.3 scientific state and 65-path Galaxy benchmark had been frozen. A new prospective external evaluation selected 1,3-propanediol and 1,4-butanediol before pathway truth was accessed. The selection record explicitly declared that evaluation truth had not been defined or accessed and that pathway literature was not used for target selection. Phase A locked generator outputs and provenance before Phase B truth comparison. DORAnet and RetroBioCat2 were evaluated as Candidate generators; backend-internal status labels were retained but were not treated as external validation. External recovery required exact-structure recovery of predeclared literature intermediates. No development ranking policy was applied in production (ranking policy NONE). The spent development/validation targets and the frozen 2.3 benchmark were not used for policy selection after truth reveal.

## 3 Results

### 3.1 Graph union preserved constituent routes and increased route diversity

Across the 24-target breadth panel, DORAnet produced candidates for all 24 targets, whereas RetroBioCat2 and RetroPath each produced candidates for 19. At least one constituent backend recovered complete routes for lactic acid, 3-hydroxypropionic acid, 1,4-butanediol and putrescine.

The ensemble preserved route-positive coverage for all four. It returned 23 routes for lactic acid, reached the 100-route cap for 3-hydroxypropionic acid, and returned one and two routes for 1,4-butanediol and putrescine, respectively. The panel therefore demonstrated route preservation and, for some targets, substantially greater route diversity, but no target was solved exclusively by cross-engine composition.

The offline route-preservation audit found zero graph-monotonicity violations, zero missing constituent reaction edges after union and zero audited constituent routes absent from ensemble enumeration. This establishes preservation under the tested normalization and enumeration settings while keeping route coverage distinct from ranking quality.

### 3.2 Held-out literature recovery broadened, but exact-route ranking did not improve

On the frozen 65-path held-out benchmark, strict stereochemical matching produced no complete literature-pathway recovery. Connectivity-level evaluation was more informative. RetroPath exactly recovered p-hydroxystyrene and phenol at rank 12, whereas RetroBioCat2 produced no complete exact pathway. Partial connectivity-level recovery was observed for 13/65 pathways with RetroBioCat2, 9/65 with RetroPath and 19/65 with the ensemble; DORAnet showed no literature-reaction overlap under the benchmark normalization.

**Figure 2.**
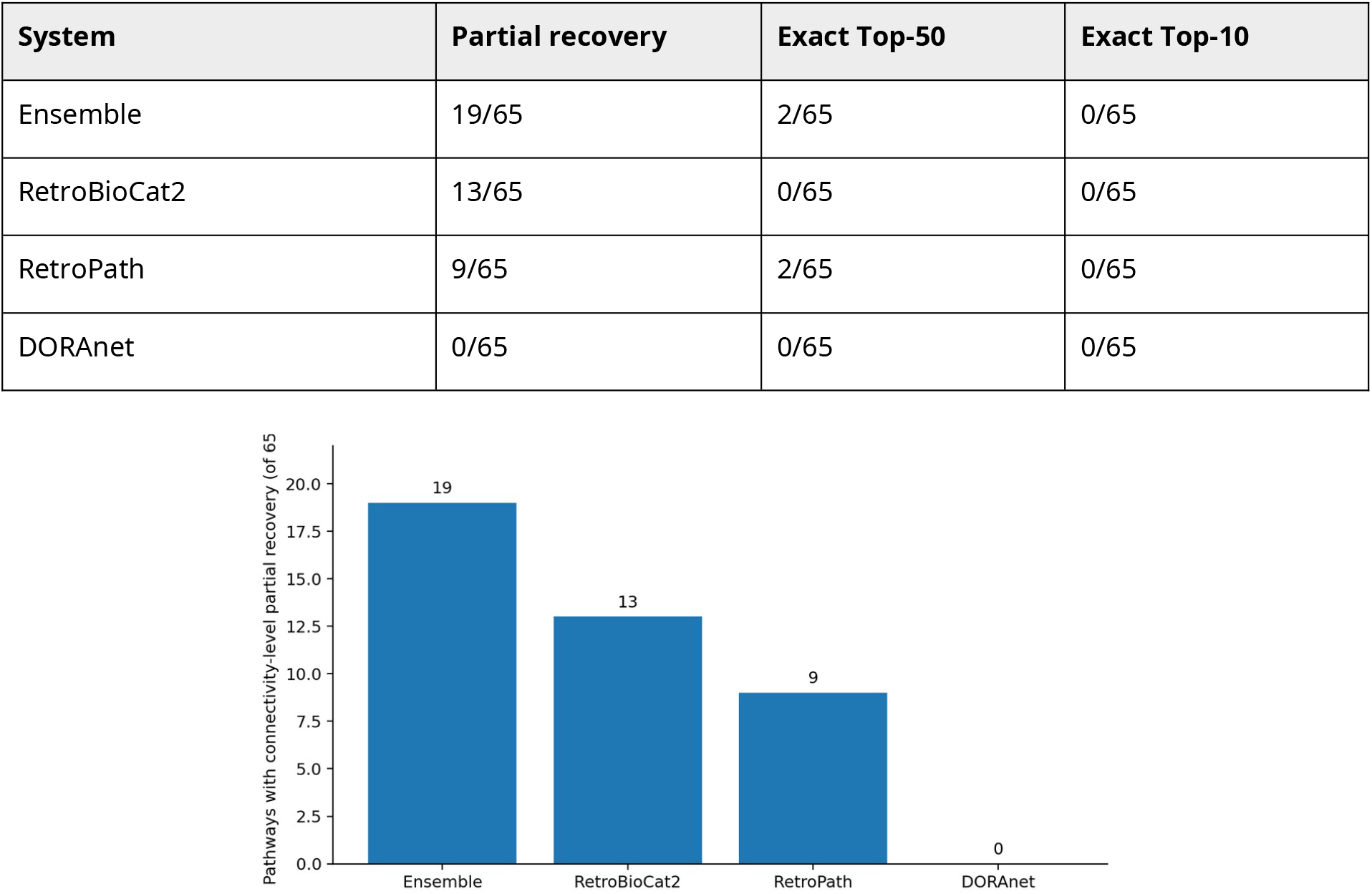
**Connectivity-level partial pathway recovery on the frozen 65-path Galaxy literature benchmark. The denominator includes one mapping-limited target; 64 targets were runnable.** *Alt text: Bar chart comparing connectivity-level partial pathway recovery on 65 held-out literature pathways: ensemble 19, RetroBioCat2 13, RetroPath 9 and DORAnet 0*.

The ensemble preserved both exact RetroPath routes, but the frozen 2D-primary ranking moved p-hydroxystyrene from rank 12 to rank 46 and phenol from rank 12 to rank 31. Ensemble exact-route MRR was therefore 0.027 versus 0.083 for RetroPath. No exact route appeared in the ensemble Top-10. Thus, the validated ensemble benefit in 2.3 is broader aggregate reaction recovery and route preservation, not superior exact Top-K ranking.

### 3.3 Failure localization identified biochemical discrimination as the dominant bottleneck

A non-tuning post-hoc classification localized the principal held-out failure mode. Of 65 pathways, 42 produced complete predicted routes but no reaction-level overlap with the reported literature pathway, 17 showed partial recovery, two achieved exact connectivity-level recovery, three were affected by backend runtime failures and one was mapping-limited. Most failures therefore occurred after candidate generation had already produced chemically complete routes. This shifts the next research problem from search-space expansion toward biochemical discrimination, evidence integration and route ranking.

**Figure 3.**
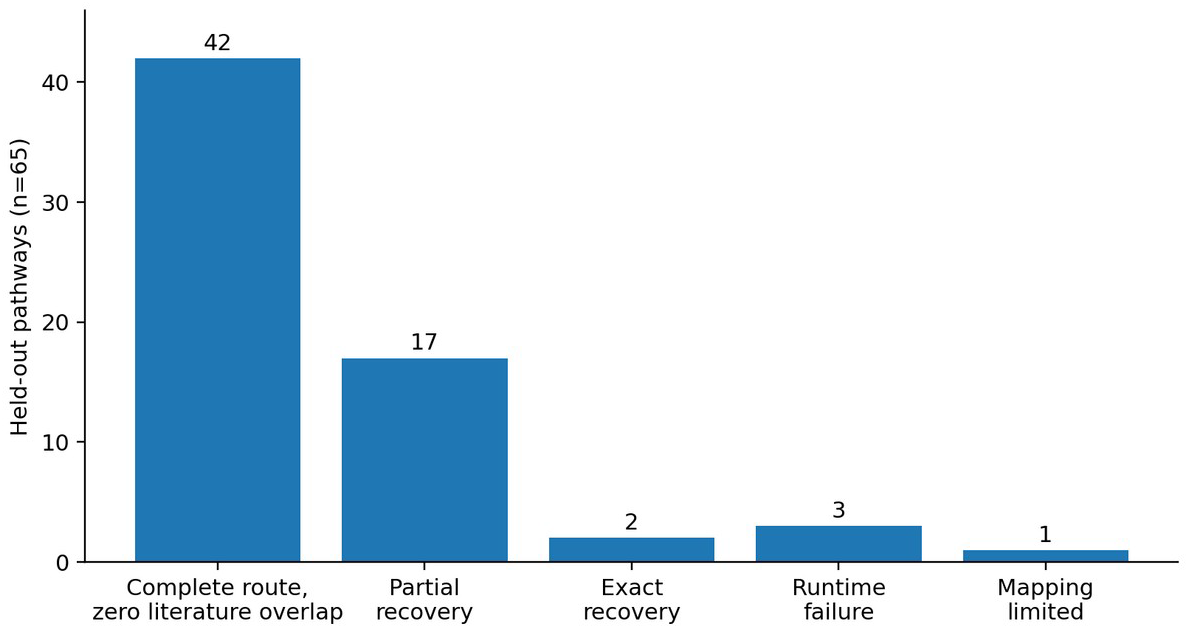
**Held-out failure classification. Forty-two of 65 targets had complete predicted routes but zero literature-reaction overlap, dominating runtime and mapping failures.** *Alt text: Bar chart of held-out failure classes: 42 complete predicted routes with zero literature-reaction overlap, 17 partial recoveries, 2 exact connectivity-level recoveries, 3 backend runtime failures and 1 mapping-limited target*.

### 3.4 Development-set ablation favored 2D-primary similarity

In leave-one-pathway-out development analysis, 2D-only ranking achieved exact Top-1 recovery for 1/12 targets, exact Top-5 recovery for 2/12 and exact-route MRR 0.750. Both 3D-first and 3D-only achieved 1/12 Top-1, 1/12 Top-5 and MRR 0.529. EC-neighbor retrieval likewise slightly favored 2D-only (Top-1 9/10, MRR 0.925) over the 3D policies (8/10, MRR 0.900). 3D-first changed the top-ranked route on 10/12 targets, showing that the signal materially altered prioritization without improving validated recovery. SynBioCrow 2.3 therefore froze 2D similarity as primary and retained 3D only as an optional secondary feature.

| Policy | Exact Top-1 | Exact Top-5 | Exact-route MRR | EC-neighbor Top-1 |
| --- | --- | --- | --- | --- |
| 2D Morgan/Tanimoto | 1/12 | 2/12 | 0.750 | 9/10 |
| 3D-first USRCAT | 1/12 | 1/12 | 0.529 | 8/10 |
| 3D-only USRCAT | 1/12 | 1/12 | 0.529 | 8/10 |

### 3.5 Evidence-aware DBTL remained conservative

A persisted 1,4-butanediol route demonstrated separation of pathway proposal from evidence. Two of four reactions passed explicit stoichiometric closure; no route edge had exact promoted Rhea support, exact reviewed-enzyme support or quantitative thermodynamic support, and the route-level evidence decision remained FAIL. Build abstained because the benchmark artifact lacked provenance-backed protein, CDS and regulatory-part sequences.

A separate sequence-backed sabinene example resolved reviewed sabinene synthase A6XH06 from Salvia pomifera, generated a 1,746-nt E. coli-preferred CDS with exact translation preservation, and removed one BamHI motif by a deterministic synonymous change while retaining 54.58% GC. Full cassette assembly abstained because verified promoter, RBS and terminator sequences were not supplied. For 20 alternative 3-hydroxypropionic-acid routes, bounded Learn changed lower-rank ordering while leaving the highest-ranked route unchanged and could not alter evidence or lifecycle state.

### 3.6 Prospective 2.4 evaluation did not recover predeclared external pathway structures

For 1,3-propanediol, DORAnet generated 30 candidates and no internally solved route, whereas RetroBioCat2 generated 1,373 candidates, including 3 backend-internal SOLVED search outcomes. For 1,4-butanediol, DORAnet generated 33 candidates and no internally solved route, whereas RetroBioCat2 generated 1,466 candidates, including 2 backend-internal SOLVED search outcomes. These SOLVED labels describe backend search termination only. Under the predeclared external criterion, neither DORAnet nor RetroBioCat2 recovered any explicit literature intermediate by exact structure for either target. The prospective external truth-structure recovery result was therefore negative for both targets. The result was frozen without post-reveal tuning, no ranking policy was promoted, and SynBioCrow 2.4 remained Candidate-stage.

## 4 Discussion

The combined 2.3 and 2.4 evidence separates three outcomes that are often conflated: broader reaction recovery, greater route diversity and externally validated recovery of complete pathways. The frozen 2.3 benchmark supports the first two but not the third: graph union preserved every audited constituent route and increased partial literature-reaction recovery from the strongest constituent value of 13/65 to 19/65, while no cross-engine-only literature route was recovered within the bounded search and the ensemble worsened the ranks of the two exact routes supplied by RetroPath. The prospective 2.4 experiment strengthens the negative conclusion: repaired Candidate generation produced substantial search output, including five RetroBioCat2 backend-internal SOLVED outcomes across two new targets, but recovered none of the predeclared external literature intermediates by exact structure.

This distinction changes the interpretation of ensembling. The useful property is not simply a larger candidate set. Provenance-preserving union makes complementary chemistry available downstream without erasing constituent solutions, but biochemical discrimination and multistep reconstruction must distinguish useful hypotheses from search volume. The 2.3 held-out error profile already showed that 42/65 targets had complete predicted routes with zero literature-reaction overlap. The 2.4 prospective result independently demonstrates why backend-internal completion cannot be equated with external pathway recovery.

The 2.3 similarity ablation provides a second negative but actionable result. USRCAT 3D-first ranking materially changed route ordering yet did not improve validated development-set retrieval over Morgan/Tanimoto 2D. That 2D-primary policy remains part of the frozen 2.3 baseline, but it was not promoted as a validated production policy for 2.4. Because subsequent development did not establish a non-circular ranking policy under prospective evaluation, SynBioCrow 2.4 freezes production ranking as NONE. This avoids converting retrospective ranking diagnostics into a prospective performance claim.

The DBTL demonstrations show how conservative evidence handling can coexist with automated design. Missing sequence or regulatory-part provenance causes abstention rather than fabricated completion, and bounded learning changes prioritization rather than evidence state. These properties are important for computational design systems intended to support, rather than replace, experimental judgment.

The next research objective is therefore improved participant/role semantics, multistep reconstruction and biochemical discrimination under a newly predeclared evaluation design. The 65-path 2.3 benchmark and the 2.4 prospective targets are now spent evidence and should not become tuning sets. Any future ranking policy should be selected without access to its evaluation truth and then tested prospectively on entirely new targets.

## 5 Limitations

All results are computational and do not establish pathway activity, enzyme productivity, titer, yield or construct function. Benchmark outcomes depend on compound normalization, precursor/sink definitions, search-depth and state caps, and the chemistry represented by each backend. The Galaxy benchmark rewards recovery of reported pathways; a chemically plausible alternative route can score as zero overlap. Conversely, connectivity-level matching intentionally ignores stereochemical distinctions and is less stringent than strict reaction identity. One 2.3 held-out pathway was mapping-limited and three were affected by backend runtime failures. The 2.4 prospective evaluation used only two new external targets and exact recovery of predeclared explicit intermediates; its negative result does not prove that generated alternatives are chemically impossible, but it does preclude a claim of validated literature-pathway recovery. Backend-internal SOLVED status is not external validation. BioPKS/RetroTide post-PKS continuation remained computationally expensive in the tested examples, and the sequence-backed sabinene example validates digital provenance and QC rather than experimental function.

## 6 Data and code availability

SynBioCrow 2.4.0 is publicly released as GitHub tag v2.4.0 and permanently archived on Zenodo at DOI 10.5281/zenodo.23173173. The validated release tag points to repository commit e2b5b6dc783c2c44b45365a5df424df95fad6704. The release supporting-data ZIP has SHA-256 9ff0f618bc5dec1047302a2d516bfb47e82b875a3ebf68b71b2351c38cd1758a. The archive contains the v2.4.0 source snapshot and frozen supporting evidence, including the prospective-evaluation record and scientific freeze material. The SynBioCrow 2.3 scientific baseline remains independently frozen and archived at DOI 10.5281/zenodo.23083426; it was not retroactively altered or reused as a 2.4 tuning set.

## 7 Author contributions

Thomas J. Heiman: Conceptualization, Methodology, Software, Validation, Formal analysis, Investigation, Data curation, Visualization, Writing - original draft, Writing - review and editing, Project administration.

## 8 Artificial intelligence assistance disclosure

OpenAI ChatGPT was used during software development for coding assistance, debugging, workflow planning, release-engineering support, and assistance with computational data-processing and analysis workflows, and during manuscript preparation for editorial restructuring, generation of draft language, and language editing. The human author defined the scientific questions and evaluation constraints, directed and executed the computational analyses, inspected outputs, reviewed AI-generated code and text before use, verified reported numerical results against frozen artifacts and primary sources, and accepts responsibility for all claims and the final manuscript. Generative AI was not treated as an author, benchmark truth, database evidence, experimental evidence, or an independent source of scientific conclusions; AI-assisted outputs were not accepted as reported results without human verification. This use is disclosed for editorial assessment under the journal’s AI policy.

